# Multiple Allosteric Sites Allow for Synergistic Enhancement of GPCR Signaling

**DOI:** 10.1101/2025.10.21.683604

**Authors:** Kai Wang, Snezana T. Dimova, John N. Hanson, Abhishek Thakur, Youwei Xu, Yini Liu, Amy E. Moritz, Alexander Goldberg, Bing Xie, Jayachandra Rayadurgam, Feijun Wang, Ashley N. Nilson, Kathryn D. Luderman, R. Benjamin Free, Wen Hu, Jian Zhang, H. Eric Xu, Kevin J. Frankowski, Yue Wang, Lei Shi, David R. Sibley, Youwen Zhuang

**Author notes:** Correspondence to: K.J.F., Y.W., L.S., D.R.S, Y.Z. These authors contributed equally to this paper (Kai Wang, Snezana T. Dimova, John N. Hanson, Abhishek Thakur, Youwei Xu).

## Abstract

Allosteric modulation of G protein-coupled receptors (GPCRs) is an emerging therapeutic paradigm that has proven effective, yet the cooperative action of multiple modulators remains unexplored. Here, we reveal how positive allosteric modulators (PAMs) synergistically enable extraordinary signal amplification through the D_1_ dopamine receptor (D1R). We developed UNC9815 and UNC10062 as enhanced D1R PAMs from the parent compound MLS6585, and then employed cryo-electron microscopy to reveal concurrent occupancy of three distinct allosteric sites by PAMs of different scaffolds, including LY3154207, BMS-A1, and our UNC compounds. Remarkably, we discovered two adjacent allosteric pockets at the transmembrane helix (TM) 1-7 interface: BMS-A1 occupies an intracellular site promoting activation through TM7 conformational shifts, while the UNC compounds complementarily stabilize the extracellular side of the interface. When combined with LY3154207, this cooperative architecture enhances dopamine potency by more than 1,000-fold. These findings elucidate the first structural blueprint for multi-site GPCR cooperativity, unlocking transformative therapeutic strategies inaccessible to orthosteric and single-site allosteric drugs.

## Introduction

G protein-coupled receptors (GPCRs) constitute the largest class of membrane receptor drug targets and mediate diverse physiological processes^1,2^. Among them, dopamine receptors (DRs) play essential roles in dopaminergic neurotransmission and signaling^3,4^. These receptors are classified into two subfamilies: D1-like receptors include the D_1_ dopamine receptor (D1R) and D5R that couple to G_s/olf_ proteins to stimulate adenylyl cyclase and increase cAMP production, whereas D2-like receptors (D2R, D3R, and D4R) activate G_i/o_ proteins to inhibit cAMP production^3,4^. The D1R, in particular, represents a critical therapeutic target due to its abundance in brain regions controlling movement, motivation, and cognition^3–5^. The role of D1R in neuropsychiatric disorders, including Parkinson’s disease (PD), schizophrenia, and Alzheimer’s disease (AD), has established it as a major focus for drug discovery^3,4,6,7^.

Traditional drug development for GPCRs have predominantly focused on the orthosteric binding pocket (OBP), where endogenous agonists bind^6,8^. However, this approach has encountered significant hurdles in the development of D1R-targeted drugs^6,8^. In conditions like PD, where dopaminergic neuron degeneration leads to reduced striatal dopamine (DA) levels, orthosteric agonists of the D1R have been explored as therapeutic alternatives to L-DOPA^3,9^. Unfortunately, their usage has been hindered by poor oral bioavailability, limited blood-brain barrier penetration, and insufficient subtype selectivity. These liabilities lead to adverse side effects including hypotension and tachyphylaxis, resulting in limited clinical applications centered around non-CNS indications^6,10–12^.

To address these challenges, allosteric modulation has emerged as an innovative strategy to overcome such limitations in GPCR drug discovery^13–15^. Unlike orthosteric ligands, allosteric modulators bind to distinct, less conserved receptor sites, modulating responses to endogenous ligands without competing for the OBP^6,14,16,17^. Compared to the orthosteric agonists, positive allosteric modulators (PAMs) of the D1R have improved receptor subtype selectivity, saturable effects that mitigate the risk of overstimulation, and preservation of the spatiotemporal dynamics of DA signaling^6,18-21^. These characteristics make D1R PAMs a promising therapeutic strategy for conditions such as PD and schizophrenia^17,22^. Significant progress has been made in discovering D1R PAMs through high-throughput screening (HTS)^6,23–25^. Notably, Bristol Myers Squibb, Eli Lilly, and Astellas have identified PAMs such as BMS-A1, LY3154207, and ASP4345, some of which have advanced to preclinical or clinical evaluation^6,26–31^. Our previous HTS work identified additional promising candidates, including MLS1082 and MLS6585^24^. MLS1082 was found to target the same intracellular loop 2 (ICL2) binding site as LY3154207, whereas MLS6585 appeared to interact with a distinct site on the D1R^23,32^. Despite these advances, most PAM candidates face translational challenges, including species-specific activity, suboptimal pharmacokinetics, and limited understanding of their binding mechanisms^6,33^.

D1R PAMs such as MLS6585, BMS-A1, and LY3154207 have been proposed to bind to distinct allosteric pockets and have shown cooperative effects when used in combination^33^. This observation reveals a new dimension of GPCR regulation and offers exceptional opportunities for fine-tuning receptor function for therapeutic development. However, the molecular basis of this cooperativity remains unclear, with only the binding site of LY3154207 definitively characterized to date^34^. To address these critical gaps, and establish a structural and mechanistic framework for cooperative allosteric modulation of the D1R, we first optimized MLS6585 through medicinal chemistry, yielding two analogs, UNC10062 and UNC9815, with enhanced potency and efficacy^35^. By combining pharmacological characterization with cryo-electron microscopy (cryo-EM) structural analyses, we identified two novel allosteric sites on the D1R that selectively interact with either BMS-A1 or the UNC compounds. Structural analyses and molecular dynamics (MD) simulations further uncovered mechanisms of synergistic allosteric regulation when multiple PAMs occupy the receptor. Together, we show how discrete modulators work together with dopamine to produce extraordinary synergistic enhancement of receptor activation.

## Results

### Mutual cooperativity of three distinct D1R PAM scaffolds

We previously described the D1R-G_s_ complex structures bound to the orthosteric agonists DA or SKF81297 together with the PAM LY3154207^34^. We had also attempted to obtain a D1R structure using these orthosteric agonists and the PAM MLS6585, but the MLS6585 binding density was not detectable, potentially due to its low binding affinity^24,34^. In a separate line of investigation (to be described elsewhere), we performed chemical optimization of MLS6585 to increase its potency and efficacy for potentiating DA affinity and signaling at the D1R, yielding optimized analogs UNC9815 and UNC10062 (Figs. 1a-b and Extended Data Figs. 1-2). We then used UNC9815 and UNC10062 to investigate the cooperative effects with LY3154207 and BMS-A1 using D1R-mediated β-arrestin recruitment assays.

**Fig. 1.**
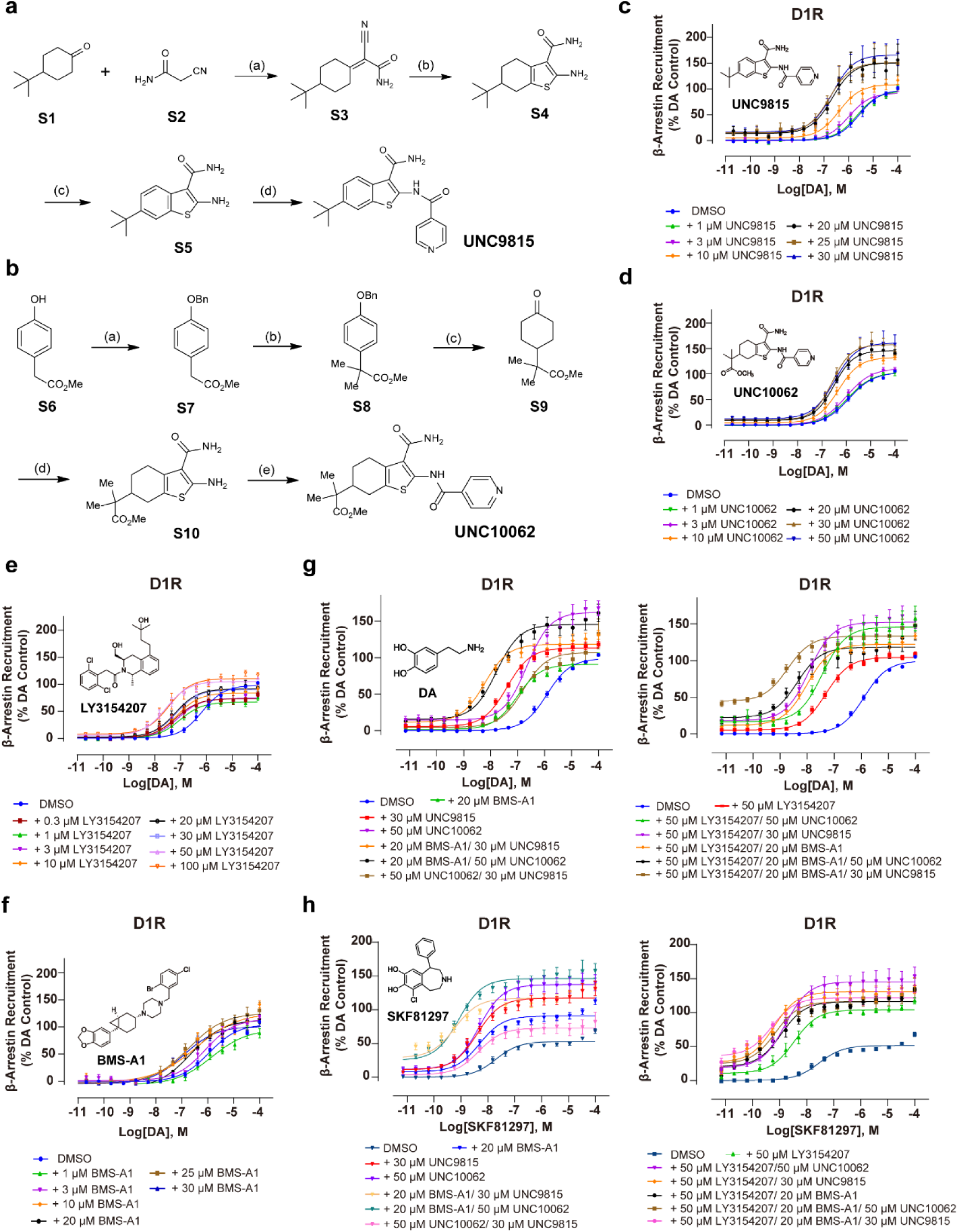
Characterization of the cooperative effects of D1R PAMs. **a,** Synthetic route for UNC9815. Reagents and conditions: (a) Ammonium acetate (1.0 equiv), trifluoroacetic acid (1.0 equiv), toluene, reflux, 67% yield; (b) sulfur (1.1 equiv), morpholine (1.0 equiv), ethanol, 50 °C, 5 h, 90% yield; (c) chloranil (2.0 equiv), 1,4- dioxane, reflux, 17 h, 71% yield; (d) isonicotinic acid (1.0 equiv), oxalyl chloride (1.5 equiv), dimethylformamide (catalytic), pyridine (2.0 equiv), dichloromethane, room temperature (RT), 2 h, 69% yield. **b,** Synthetic route for UNC10062. Reagents and conditions: (a) benzyl bromide (1.0 equiv), potassium carbonate (3.0 equiv), DMF, RT, 85% yield; (b) sodium hydride (3.0 equiv), methyl iodide (3.0 equiv), THF, 0 °C to RT, 29% yield; (c) Pd/Al_2_O_3_ (5 wt%), hydrogen (5 bar), dichloroethane, 54% yield; (d) sulfur (1.0 equiv), 2-cyanoacetamide (1.5 equiv), morpholine (1.5 equiv), ethanol, 120 °C, microwave irradiation, 0.5 h, 60% yield; (e) isonicotinoyl chloride hydrochloride (1.5 equiv), triethylamine (3.0 equiv), dichloromethane, 0 °C, 4 h, 32% yield. The procedures for synthesizing UNC9815 and UNC10062 are described in the **Materials and Methods. c-f,** UNC9815, UNC10062, LY3154207, and BMS-A1 dose- dependently increase DA’s potency in a saturable manner as measured in β-arrestin recruitment assays. Increasing concentrations of UNC9815, UNC10062, BMS-A1, or LY3154207 were used to potentiate DA-stimulated β-arrestin recruitment using the DiscoverX assay as described in the Methods Section. Data are displayed as a percentage of the maximum DA control response in each experiment (mean ± SEM) of ≥ 3 experiments, each performed in triplicate. Curve parameters are shown in **Extended Data Table 1. c**, UNC9815 maximally increased DA potency at 30 µM, with an EC_50_ fold-shift of 17.1 ± 2.2**. **d,** UNC10062 maximally increased DA potency at 50 µM, with an EC_50_ fold-shift of 3.9 ± 0.5**. **e,** LY3154207 (LY) maximally increased DA potency at 50 µM, with an EC_50_ fold-shift of 27 ± 4.6* **f,** BMS-A1 maximally increased DA potency at 20 µM, with an EC_50_ fold-shift of 6.3 ± 0.4**. The PAM EC_50_ fold-shift statistical comparisons were made to the control DA EC_50_ fold-shift values using a one- way ANOVA with Dunnett’s post hoc comparison: *p < 0.05, **p < 0.01. **g-h,** Cooperative effects of UNC compounds (UNC9815 and UNC10062), LY3154207, and BMS-A1 on potentiating DA- or SKF81297-induced D1R activation. β-arrestin recruitment was measured using the DiscoverX assay. Data are displayed as a percentage of the maximum control DA response in each experiment and represent mean ± SEM values from either ≥ 9 (**g**) or ≥ 6 (**h**) experiments, each performed in triplicate. Curve parameters and statistical analyses are displayed in **Extended Data Table 2**.

Initially, we performed curve-shift assays to establish a maximally effective dose of each PAM for increasing the potency of DA for stimulating β-arrestin recruitment to the D1R. As shown in Fig. 1 and Extended Data Table 1, each PAM dose-dependently increased the functional potency of DA (decrease in EC_50_) in a saturable manner while their effects on DA efficacy (Emax) were more modest, although the UNC compounds appeared to be more active in this regard (Figs. 1c-f). PAMs can promote receptor signaling either through increasing agonist potency (EC_50_), agonist efficacy (Emax) or through direct allosteric agonism, or a combination of these effects. Notably, the effects of PAMs on GPCR signaling can be remarkably assay-dependent^14^. Thus, for consistency and comparison across multiple assays in this study, we have operationally defined the PAM-induced increase in agonist potency (EC_50_) as the most reliable and quantifiable indicator of allosteric efficacy while still reporting effects on Emax and allosteric agonism. Using the EC_50_ fold-shift results in Fig. 1 and Extended Data Table 1, the D1R PAMs exhibited the following rank order of activity: LY3154207 <u>∼</u> UNC9815 > BMS-A1 > UNC10062.

Having established maximally effective concentrations for each D1R PAM, we performed additivity experiments and found that combinations of these PAMs synergistically increased the functional potencies of both DA and SKF81297, inducing ≥ 1,000-fold shift in the DA EC_50_ when the three divergent classes of PAMs were combined with DA (Figs. 1g-h and Extended Data Table 2). Interestingly, using SKF81297, a partial agonist in this assay, the three PAMs were functionally additive with each other; however, the triple PAM combination did not potentiate the SKF81297 EC_50_ beyond that observed with any double PAM combination (Figs. 1g-h and Extended Data Table 2). This observation suggests a ceiling effect such that the D1R is maximally responsive to SKF81297 when bound to at least two of the PAMs. Regardless, these findings reveal a complex allosteric modulation of the D1R, and importantly, that three distinct allosteric binding sites must exist on the D1R, which might be leveraged to synergistically enhance receptor function (Fig. 2a).

**Fig. 2.**
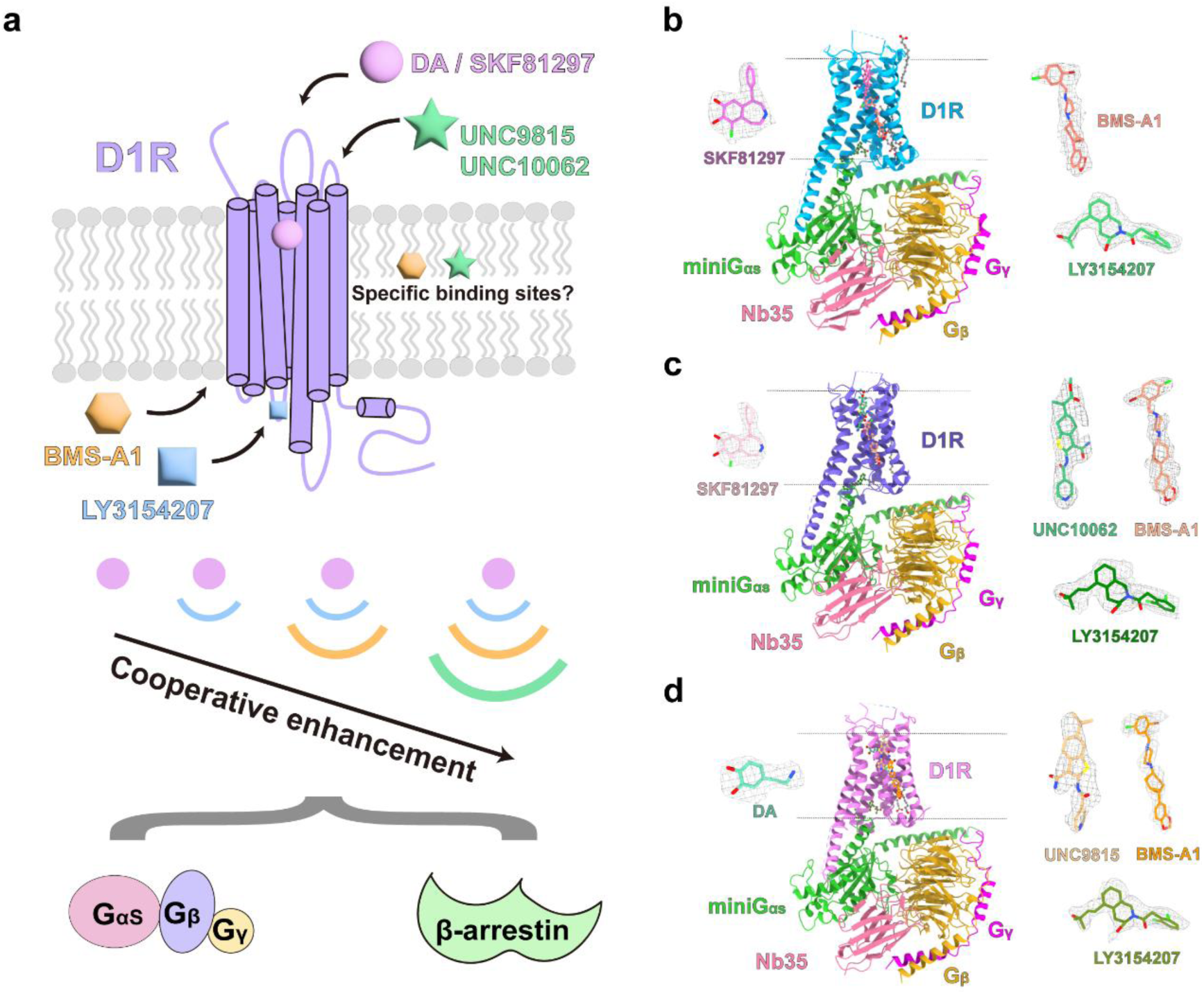
Structural basis of D1R recognition by divergent PAMs. **a,** Schematic illustration of synergistic signaling modulation by orthosteric agonists and PAMs at the D1R. Binding of DA or SKF81297 to the orthosteric pocket of D1R enables synergistic enhancement of downstream G protein and β-arrestin signaling in the presence of PAMs LY3154207, BMS-A1, and UNC9815/UNC10062. The binding pockets of DA/SKF81297 and LY3154207 are established, whereas the specific sites for BMS-A1 and UNC9815/UNC10062 remain undefined. **b,** Model of the D1R-G_s_ complex bound to orthosteric agonist SKF81297 (orchid), and allosteric modulators LY3154207 (forest green) and BMS-A1 (salmon). **c,** Model of the quaternary complex containing D1R-G_s_ bound to SKF81297, LY3154207, BMS-A1, and UNC10062 (medium sea green). **d,** Model of the D1R-G_s_ complex in complex with DA (medium aquamarine), LY3154207, BMS-A1, and UNC9815 (burly wood). The cryo-EM densities of the ligands are shown at contour levels of 0.35 **(b)**, 0.25 **(c)**, and 0.25 **(d)**, respectively. The contour level of BMS-A1 density in the DA/LY3154207/BMS-A1/UNC9815 quaternary complex is shown at 0.1.

### Structures of D1R bound to divergent PAM combinations

To comprehensively elucidate the molecular interaction and mechanism of these allosteric modulators, including BMS-A1, UNC9815, and UNC10062, we determined the structures of D1R-G_s_ signaling complexes bound to these PAMs. Using the potent agonist SKF81297 to occupy and stabilize the OBP of D1R^34,36^, we focused first on determining the structure of the D1R in complex with BMS-A1 alone or BMS-A1 and UNC10062 together. As our functional assays demonstrated that SKF81297 is a partial agonist compared to DA in the β-arrestin recruitment assay (Figs. 1g-h and Extended Data Table 2), we also used DA to occupy the OBP to investigate the PAM effects on full agonist binding with cryo-EM analysis. Notably, since LY3154207 binds to a divergent site as these PAMs and exerts synergistic PAM effects^33^, to better stabilize the protein complexes for structure determination, we incorporated LY3154207 into all sample preparation processes.

The cryo-EM structures of D1R-G_s_ in complex with SKF81297/LY3154207/BMS-A1, SKF81297/LY3154207/BMS-A1/UNC10062, and DA/LY3154207/BMS-A1/UNC9815 were determined at global resolutions of 2.65 Å, 2.59 Å, and 2.44 Å, respectively (Fig. 2b-d, Extended Data Figs. 3-4, and Extended Data Table 3). The high-resolution maps allowed us to unambiguously model most portions of the D1R-G_s_ complex, the orthosteric agonists SKF81297 or DA, and PAMs. As previously reported^34,37–39^, LY3154207 bound above the ICL2 within the D1R. In the map of D1R- SKF81297/LY3154207/BMS-A1, we observed robust density of BMS-A1 at the intracellular side of the receptor with BMS-A1 sandwiched between TM1 and TM7 in a vertical conformation above Helix 8 (H8) (Fig. 2b and Extended Data Fig. 4a), a positive allosteric site similar to those identified in the D2R^40^, A_1_ adenosine receptor (A_1_AR)^41^, A_3_AR^42^, and free fatty acid receptor 2 (FFAR2)^43^. While BMS-A1 density was confirmed in the other two maps, D1R-SKF81297/LY3154207/BMS- A1/UNC10062 and D1R-DA/LY3154207/BMS-A1/UNC9815, we also observed distinct densities above BMS-A1 at the extracellular portion of the interface between TM1 and TM7, attributable to the UNC compounds in vertical orientations in the corresponding maps (Figs. 2c-d and Extended Data Figs. 4b-c). UNC9815 and UNC10062, have high structural similarity and bound to the same pocket with comparable orientations. The stacked arrangement of two PAMs sandwiched between TM1 and TM7 represents, to our knowledge, a novel allosteric binding configuration previously unobserved in other GPCRs. Interestingly, in the map of D1R- SKF81297/LY3154207/BMS-A1, we observed a clear density of cholesterol at a site similar to that occupied by UNC compounds (Figs. 2b-d, and Extended Data Fig. 4-5). This intriguing observation suggests that the UNC compounds and cholesterol may play similar roles in allosterically modulating D1R function.

### Binding of D1R PAMs in divergent allosteric sites

The binding interface between the D1R and its allosteric modulators, including BMS- A1 and both UNC compounds, is predominantly characterized by hydrophobic, aromatic, and van der Waals interactions (Figs. 3a-c). In our structures, BMS-A1 occupies a large vertically extended pocket spanning from the middle to the intracellular end of TM7 and above H8 (referred to as the “lower pocket”). The lower pocket is mainly formed by residues from TM1, TM7, and H8, including F29^1.38^, L32^1.41^, L33^1.42^, S36^1.45^, G40^1.49^, S325^7.47^, L326^7.48^, I329^7.51^, F333^7.55^, and F341^8.54^ (Fig. 3a). Specifically, the benzodioxole moiety of BMS-A1 points to H8 and interacts with F333^7.55^ and F341^8.54^, while the 2-bromo-5-chlorobenzyl group stacks with F29^1.38^, L32^1.41^, and L33^1.42^ (Fig. 3a). In addition to hydrophobic interactions, the piperazine ring of BMS- A1 forms a polar hydrogen-bond interaction with S36^1.45^ (Fig. 3a).

**Fig. 3.**
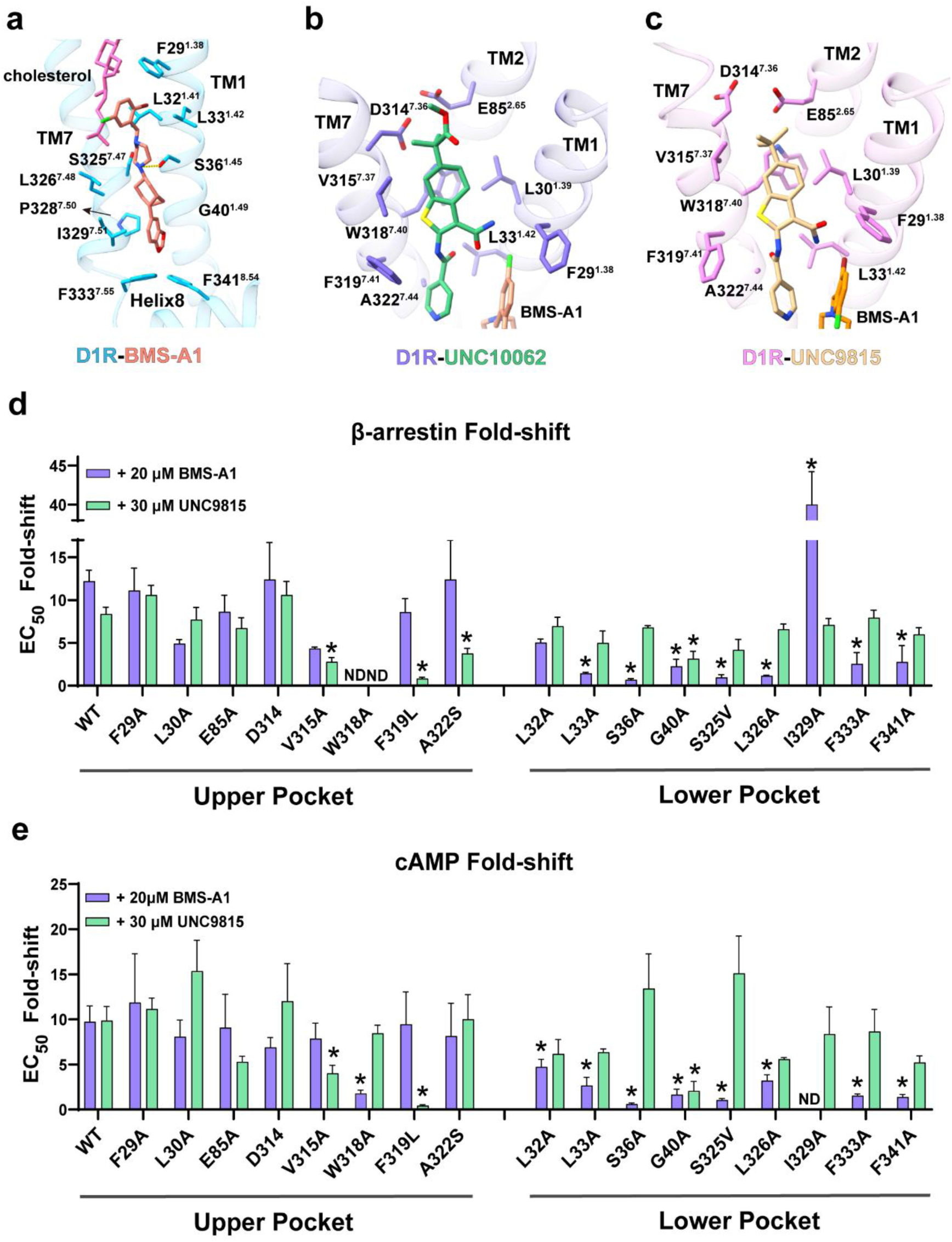
Binding interactions and effects of D1R mutations on UNC9815 and BMS- A1 PAM activities and comparison of the binding modes of D1R PAMs. **a,** Binding pose and interaction profile of BMS-A1 within the PAM pocket of the D1R-G_s_ complex containing SKF81297 and LY3154207. **b,** Binding mode and key interactions of UNC10062 within the allosteric pocket of the SKF81297/LY3154207/BMS- A1/UNC10062-bound D1R-G_s_ complex. **c,** Binding pose and interaction residues of UNC9815 in the allosteric pocket of the DA/LY3154207/BMS-A1/UNC9815-bound D1R-G_s_ complex. **d,** EC_50_ fold-shifts induced by BMS-A1 and UNC9815 in the β- arrestin BRET assays shown in **Extended Data Figs. 4 and 5 (CRCs)**, and **Extended Data Tables 4** and **5** (CRC parameters). Asterisks indicate EC_50_ fold-shift values observed with the mutant D1Rs that significantly differ from the WT D1R EC_50_ fold- shift values as shown in **Extended Data Tables 4** and **5**. ND = EC_50_ not determinable. **e,** EC_50_ fold-shifts induced by BMS-A1 and UNC9815 in the CAMYEL cAMP BRET assays shown in **Extended Data Figs. 6 and 7 (CRCs)**, and **Extended Data Tables 6** and **7** (CRC parameters). Asterisks indicate EC_50_ fold-shift values observed with the mutant D1Rs that significantly differ from the WT D1R EC_50_ fold-shift values as shown in **Extended Data Tables 6** and **7**. ND = EC_50_ not determinable.

The binding poses of both LY3154207 and BMS-A1 remains largely preserved following the addition of the UNC compounds (Extended Data Figs. 6a-b). However, the 2-bromo-5-chlorobenzyl group of BMS-A1 undergoes notable conformational changes due to specific interactions with these UNC compounds (Extended Data Fig. 6b). While both UNC compounds occupy a similar allosteric site located in the extracellular cleft between TM1 and TM7 (referred to as “upper pocket”), positioned directly above the BMS-A1 binding pocket, they exhibit distinct binding modes (Figs. 3b-c, and Extended Data Fig. 6c). Specifically, UNC9815 penetrates approximately 2 Å deeper toward the intracellular side compared to UNC10062 when measured at the α-carbon atoms of their respective carbamoyl groups (Extended Data Fig. 6c). This difference in positioning may be attributed to the more rigid planar benzothiophene core structure of UNC9815 compared to the relatively flexible half-boat conformation of the tetrahydrobenzothiophene moiety of UNC10062 (Figs. 1a-b, and 3b-c). Unlike the BMS-A1/UNC10062-bound D1R, little density of the 2-bromo-5-chlorobenzyl group of BMS-A1 could be observed in the BMS-A1/UNC9815-bound structures due to the conformational flexibility of this group (Figs. 2c-d). The hydrophobic pocket to which both UNC10062 and UNC9815 vertically bind is mainly constituted by F29^1.38^, L30^1.39^, L33^1.42^, E85^2.65^, D314^7.36^, V315^7.37^, W318^7.40^, F319^7.41^, and A322^7.44^ (Figs. 3b, c).

To confirm the above structural findings, we performed extensive mutagenesis experiments involving the PAM contact residues within the D1R TM1-TM7 upper and lower allosteric pockets. Initially, we created single residue mutants and evaluated their effects on BMS-A1 and UNC9815 PAM activities using DA as the orthosteric agonist. We chose to evaluate UNC9815 as the representative UNC D1R PAM as it is more efficacious than UNC10062 in potentiating the potency of DA (Fig. 1 and Extended Data Table 1). Further, we evaluated the D1R point mutants using both β-arrestin recruitment (Fig. 3d, Extended Data Figs. 7-8, and Extended Data Tables 4-5) and cAMP accumulation assays (Fig. 3e, Extended Data Figs. 9-10, and Extended Data Tables 6-7). Consistent with the above structural findings, mutation of most of the surrounding residues of the upper pocket, such as V315A, F319L, and A322S, attenuated the DA EC_50_ fold-shift observed with UNC9815 (Figs. 3d, e, Extended Data Figs. 7 and 9, Extended Data Tables 4 and 6). In contrast, mutation of just one lower binding pocket residue, G40A, affected UNC9815 activity, which was observed in both functional assays (Figs. 3d, e). The conserved G40^1.49^ residue had been found to introduce flexibility and bending into TM1^44^, leading to a distinct orientation of its extracellular portion compared with receptors containing P39^1.48^. Thus, the G40A mutation at the intracellular side of TM1 is expected to propagate structural effects to the extracellular end of the helix.

Mutation of most of the lower pocket residues, including L32A, L33A, S36A, G40A, S325V, L326A, I329A, F333A and F341A significantly affected the PAM activity of BMS-A1 in one or both D1R functional assays (Figs. 3d, e, Extended Data Figs. 8 and 10, Extended Data Tables 5 and 7). In most cases, these mutations attenuated the BMS- A1 PAM activity, however, in the β-arrestin recruitment assay, the I329A mutation produced a significant gain of BMS-A1 PAM efficacy (Fig. 3d and Extended Data Fig. 8, Extended Data Table 5). Only one upper pocket residue, W318A, attenuated BMS- A1 activity and only in the cAMP assay (Extended Data Figs. 8 and 10). Taken together, these experiments strongly support the poses of UNC9815 and BMS-A1 in the upper and lower pockets, respectively, as observed in the cryo-EM experiments.

To further validate the PAM binding poses revealed by the cryo-EM structures, we created D1R cluster mutants of either the upper or lower pockets or both. For these mutants, we chose residues that, when mutated, had minimal effects on DA potency yet had significant effects on PAM activity, as observed in the single mutant experiments.

As with the point mutation studies, we utilized a single UNC scaffold (UNC9815) and employed a β-arrestin recruitment assay. As shown in Extended Data Fig. 11 and Extended Data Table 8, when using the WT D1R, dual and triple combinations of UNC9815, BMS-A1 and LY3154207 exhibited additive effects on potentiating the potency of DA, resulting in a 1,500-fold leftward shift in the DA EC_50_ with all three PAMs combined. In the D1R upper cluster mutant, consisting of V315A-F319L-A322S, BMS-A1 and LY3154207 produced similar results as seen with the WT receptor (including in combination). However, UNC9815 surprisingly behaved as a negative allosteric modulator (NAM) when tested alone or in combination with BMS-A1 or LY3154207 (Extended Data Fig. 11 and Extended Data Table 8). In all cases, UNC9815 significantly shifted DA’s EC_50_ to lower potency compared to that seen in the corresponding controls. These results suggest that UNC9815 still binds to the receptor with the cluster mutations in the upper pocket, however, these mutated residues have altered the allosteric efficacy of this modulator changing it from a PAM to a NAM. In contrast, in the lower cluster D1R mutant, consisting of L326A-F333A, the PAM activity of UNC9815 was retained, as well as that of LY3154207, however, BMS-A1 was rendered functionally inactive for altering DA potency either alone or in combination with UNC9815 or LY3154207 (Extended Data Fig. 11 and Extended Data Table 8). Similarly, BMS-A1 was inactive in the D1R upper/lower cluster mutant (D1R- F319L-L326A-F333A) whereas the PAM activity of LY3154207 was retained. In contrast, UNC9815 exhibited weak, but significant, NAM activity when tested with DA alone, but was functionally inactive when tested in combination with BMS-A1 and/or LY3154207 (Extended Data Fig. 11 and Extended Data Table 8). Taken together, these results confirm the receptor interaction sites of all three PAM scaffolds and also serve to illustrate the allosteric flexibility of the D1R.

### Stable occupation of BMS-A1 and UNC compounds in D1R

To assess the stability and conformational flexibility of BMS-A1 and UNC binding modes observed in the cryo-EM structures of D1R, we performed molecular dynamics (MD) simulations on all three experimentally determined complexes (Extended Data Table 9). Unlike the cryo-EM study, which was conducted in detergent micelles, our simulations embedded D1R in an explicit 1-palmitoyl-2-oleoyl-sn-glycero-3- phosphocholine (POPC) bilayer, providing a complementary view of receptor-ligand dynamics in a more physiologically relevant lipid environment. BMS-A1 contains a piperazine ring that forms a polar hydrogen bond with S36^1.45^ in the cryo-EM structure and may carry a positive charge depending on its local environment, suggesting that this feature could influence binding. However, predicting the pKa of membrane-bound ligands remains challenging, as most algorithms rely on empirical models trained on aqueous systems and lack accuracy in lipid environments. Using Epik (Schrödinger), we predicted the protonated form of BMS-A1 to be energetically favored over the neutral form, albeit with a modest state penalty of 1.2 kcal/mol (within the typical error range of such calculations). To evaluate the impact of protonation state on binding stability, we performed MD simulations of D1R with both the charged and neutral forms of BMS-A1 (Extended Data Table 9).

Analysis of the MD simulations revealed that, contrary to the initial Epik prediction, the neutral form of BMS-A1 maintained a vertically extended pose within the lower pocket (Extended Data Fig. 12). In contrast, the charged form exhibited reduced stability, frequently adopting flipped conformations or fully dissociating from the pocket. These results suggest that the neutral state of BMS-A1 forms a more stable interaction at the lower pocket in the membrane-embedded receptor. For the bound UNC compounds, we observed subtle deviations from their cryo-EM binding poses, likely due to the distinct simulation environment; however, they largely retained their overall orientations at the upper site throughout most simulations (Extended Data Fig. 12). To assess the stability of the equilibrated binding modes of BMS-A1 and the UNC compounds during simulations, we calculated pairwise ligand root-mean-square deviation (RMSD). Both compounds consistently exhibited low RMSD values, indicating stable binding within their respective pockets between TM1 and TM7 (Extended Data Fig. 12). Interestingly, UNC9815, lacking the polar methylpropanoate group, displayed a tendency to shift between two positions within the predominantly hydrophobic upper pocket. This movement did not impact the stable binding of BMS- A1 in the lower pocket (Extended Data Fig. 12). These results suggest that the membrane environment could preserve the PAM poses observed in the cryo-EM structures.

Comparison of the binding poses of BMS-A1 and UNC compounds in the absence of each other showed that both PAMs maintained stable interactions within their respective allosteric pockets. BMS-A1 remained stably bound in the lower pocket across all simulations, regardless of UNC compound occupancy in the upper site, as indicated by low ligand RMSD values and consistent interaction profiles. However, some conformational flexibility was observed in the 2-bromo-5-chlorobenzyl moiety of BMS-A1, in the absence of the bound UNC compounds, consistent with the weaker electron density for this group in the cryo-EM structures and a slight shift in pairwise RMSD (Extended Data Fig. 12). Similarly, UNC compounds maintained a stable pose in the upper pocket, closely aligned with their experimental conformations, in the absence of BMS-A1. These results indicated that, although co-binding may influence local pocket organization, each PAM was independently capable of adopting a stable binding mode within its respective site.

### Selective determinants of BMS-A1 and UNC compounds to DRs

We have previously determined the active-state structures of all DRs^45^. Comparison of these structures with the new D1R structures bound to BMS-A1 and UNC compounds allowed us to identify the molecular determinants for receptor subtype selectivity of these PAMs. Within the DR family, the D1R exhibits the highest sequence and structural homology with D5R. Despite considerable conservation in TM1 and TM7 between D1R and D5R (Fig. 4a), a notable substitution is the replacement of S36^1.45^ in D1R by W53^1.45^ in D5R. This substitution eliminates the essential hydroxyl group required for optimal BMS-A1 binding and introduces steric hindrance that impedes BMS-A1 interaction (Fig. 4b), accounting for the selectivity of BMS-A1 to D1R when compared to D5R^33^. Consistent with these structural insights, we observed that introducing an S36W mutation in the D1R diminished BMS-A1 PAM activity, while making the reciprocal W53S mutation in D5R engendered BMS-A1 PAM activity (Figs. 4c-d, and Extended Data Table 10). The upper pocket around TM1 and TM7 in D5R remains largely conserved relative to D1R, with the exception of F29^1.38^ in D1R being replaced by L46^1.38^ in D5R (Extended Data Figs. 13a, b). Consistent with these observations, we previously demonstrated that MLS6585, the parent compound of UNC10062 and UNC9815, potentiates dopamine-stimulated β-arrestin recruitment to the D5R^24^, aligning with our current findings.

**Fig. 4.**
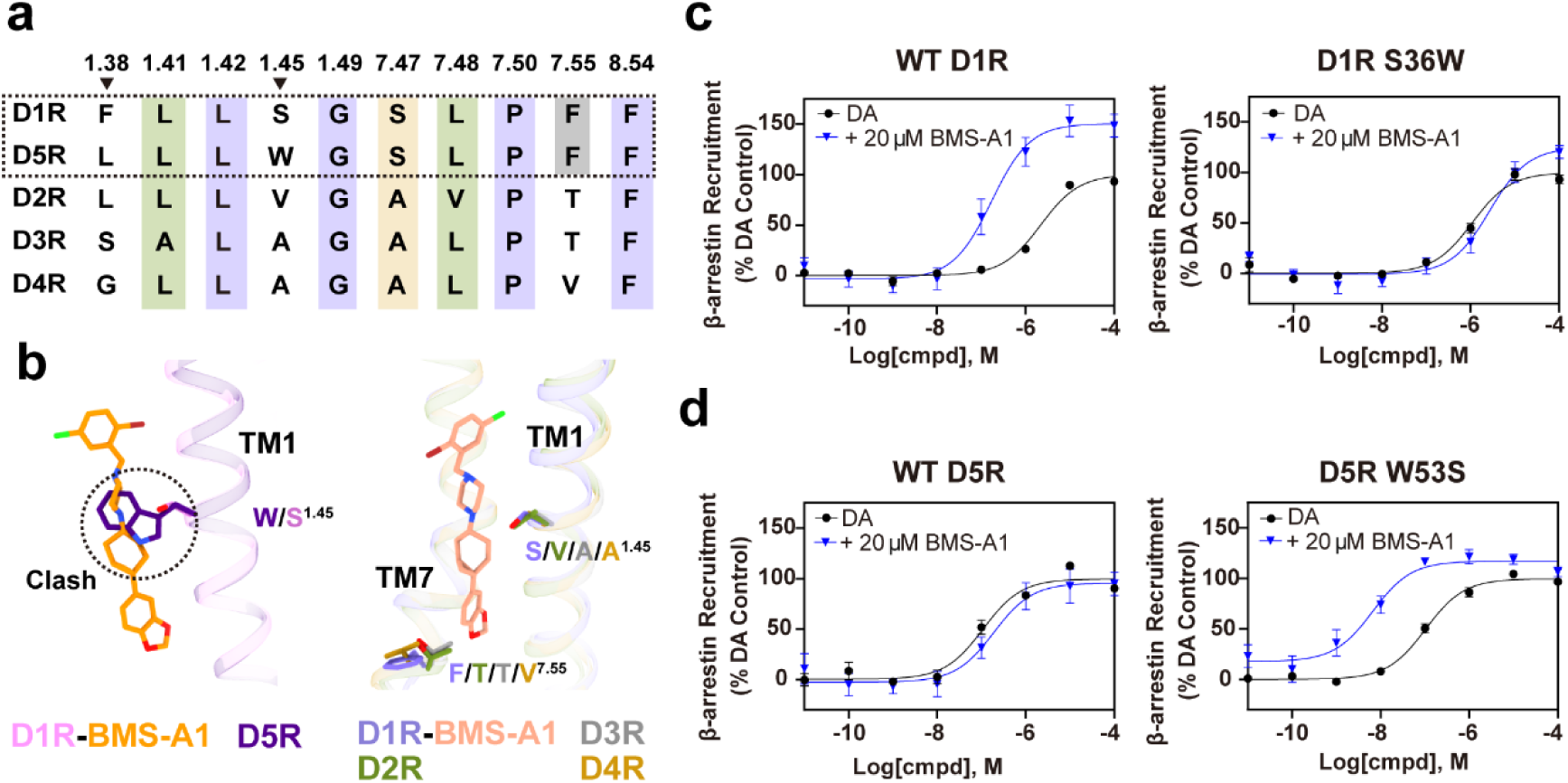
Determinants of D1R selectivity by BMS-A1. **a,** Sequence alignment of dopamine receptor subtypes highlighting PAM pocket residues critical for BMS-A1 selectivity. **b,** Structural overlay of BMS-A1-bound D1R with rotigotine-D5R (PDB: 8IRV) shows a steric clash introduced by W53^1.45^ in D5R replacing S36^1.45^ in D1R (left panel); superposition of D1R and D2-like receptors indicates non-conserved substitutions at positions 1.45 and 7.55, contributing to BMS-A1 selectivity (right panel). Rotigotine-D2R (PDB: 8IRS); rotigotine-D3R (PDB: 8IRT); rotigotine-D4R (PDB: 8IRU). **c-d,** Effects of swapping D1R and D5R residues S36^1.45^ and W53^1.45^ on BMS-A1 PAM activity using BRET β-arrestin recruitment assays as described in the Methods. Concentration-response curves of DA in the presence or absence of 20 µM BMS-A1 were generated. The data are expressed as a percentage of the control DA Emax in each experiment (n = 4) and represent mean ± SEM values. (**c**) D1R WT and S36W mutant, (**d**) D5R WT and W53S mutant. See **Extended Data Table 10** for complete curve parameters and statistical analyses.

In D2-like receptors, the residue composition of both BMS-A1 and UNC compound binding sites differs significantly from D1R, with key amino acid substitutions that create unfavorable binding conditions in D2-like receptors (Figs. 4a-b and Extended Data Fig. 13). Additionally, the conformation of the conserved W^7.40^ in D3R and D4R is rearranged by surrounding residues with bulky side chains, with its indole ring potentially introducing substantial steric constraints against UNC compounds (Extended Data Figs. 13c-f). These structural differences collectively explain the selectivity of these compounds for D1R over D2-like receptors^24^.

### Positive allosteric mechanisms of BMS-A1 and UNC compounds

Previous studies have demonstrated that LY3154207 occupies an allosteric site directly above the ICL2 in a boat-shaped configuration^34,37–39^. It exerts PAM activity through stabilization of the helical conformation of ICL2, which was then propagated to affect the conformation of the TM domain^39^. BMS-A1 and the UNC compounds, however, bind two other distinct allosteric sites, suggesting divergent mechanisms of allosteric modulation.

We recently determined the structure of D1R bound to the inverse agonist LE300 (X.Z., *et al*, in preparation). To characterize conformational changes associated with receptor activation and positive allosteric modulation, we used the Protein Interaction Analyzer (PIA)^46^ to examine three functional states of D1R: the inactive (LE300-bound), the active (agonist-bound), and the super-active (PAM-bound). Comparison of inactive and active states revealed hallmark activation changes, including F281^6.44^ moving away from the OBP center and outward displacement of TM6i, consistent with our previous observations across aminergic GPCRs^47^ (Extended Data Fig. 14). Additionally, in the active state, the extracellular portion of TM6 (TM6e, residues 6.55-6.59) moves towards, while S188^ECL2.52^ in extracellular loop 2 (ECL2) shifts away from other binding-site residues (Extended Data Fig. 14). Interestingly, PAM binding did not further displace F281^6.44^ from the OBP center but consistently induced inward movement of E85^2.65^ with either DA or SKF81297, a rearrangement likely stabilizing the super-active conformation (Extended Data Fig. 15a). Notably, S188^ECL2.52^ exhibited increased separation from binding site residues in the LY3154207/BMS-A1- and LY3154207/BMS-A1/UNC10062-bound states with SKF81297 but not with DA, while only in the triple PAM-bound state did TM6e move further inward (Extended Data Fig. 15). Thus, given that SKF81297 is a partial agonist, these ECL2 and TM6e motions proceed in consistent directions from inactive to increasingly active states.

Comparison between the inactive and SKF81297-bound active state also revealed inward movement of TM7i, which facilitates the outward tilting of TM6i, leading to opening of the G protein-binding cavity on the intracellular side of the TM domain, critical steps in signaling^48,49^. Upon BMS-A1 binding, we observed a more significant inward movement of TM7i, accompanied by notable conformational rearrangements of the conserved NPxxY motif, Y^5.58^-Y^7.53^ motif (Fig. 5a). The polar interaction between residues T^1.46^ and S^7.47^ appear to play a crucial role in maintaining the inactive state of D1R, as demonstrated in our companion study (X.Z., *et al*, in preparation), but is disrupted upon activation, driving outward movement of TM1i. BMS-A1 binding further amplifies this disruption, inducing greater outward displacement of TM1i (Fig. 5b). It is noteworthy that LY3154207 binding alone does not produce additional TM1i movement beyond the SKF81297-bound state, indicating that LY3154207 and BMS- A1 exert distinct allosteric effects on D1R. Furthermore, binding of UNC compounds to the upper allosteric site potentially stabilizes the TM7 extracellular region in its active conformation (Extended Data Fig. 16). Together, these mechanisms may underlie the synergistic increase in PAM activity observed with different combinations of LY3154207, BMS-A1, and UNC compounds.

**Fig. 5.**
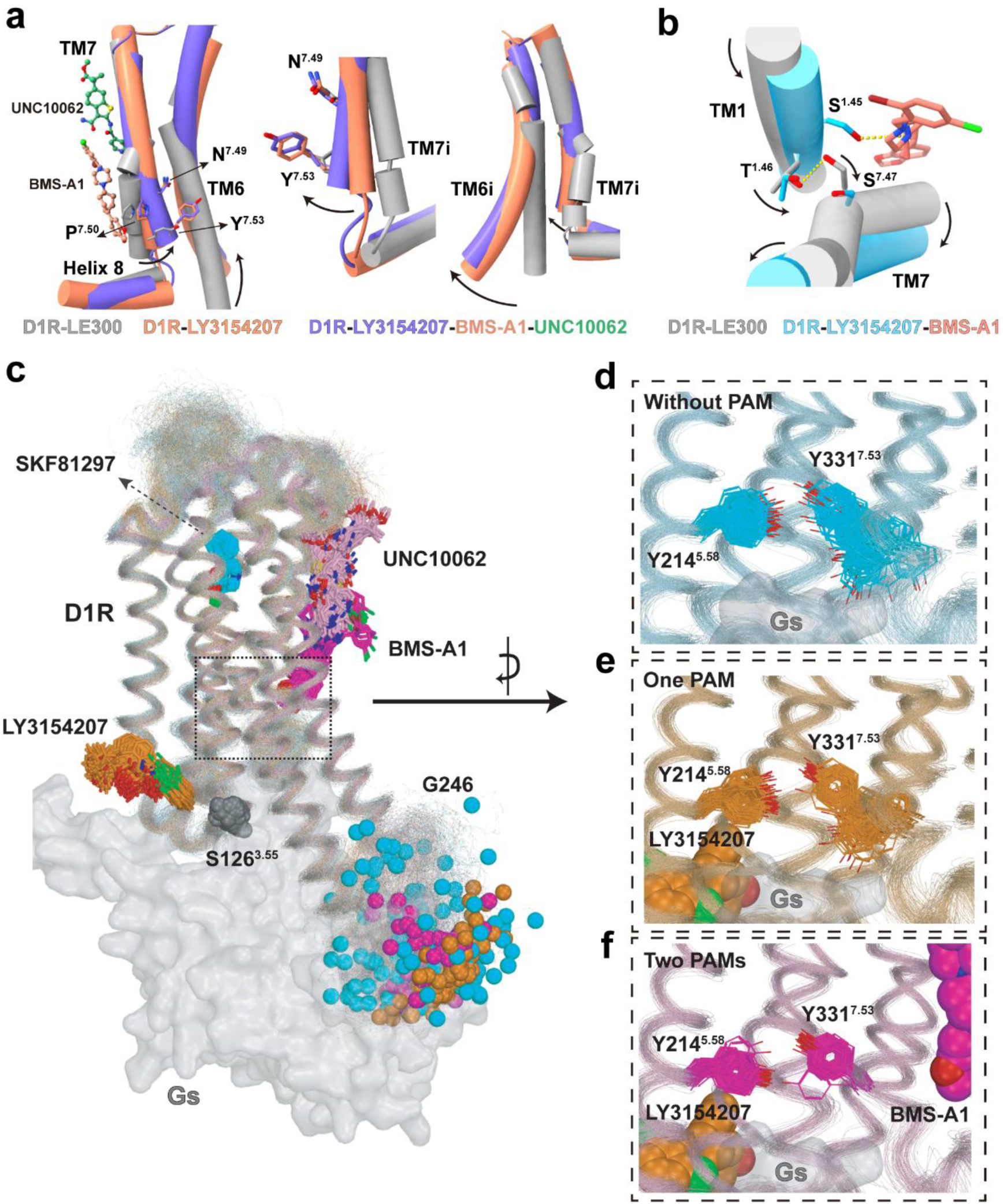
Conformational changes in D1R induced by PAM binding. **a,** Structural alignment of D1R in inactive, active, and super-active (PAM-bound) states highlights inward movement of TM7i and outward movement of TM6i upon activation. **b,** Top- down view of D1R illustrating disruption of the T^1.46^-S^7.35^ hydrogen bond upon BMS- A1 binding. **c,** The overall similar receptor dynamics across three comparative simulations conditions: D1R-SKF81297 (cyan), D1R-SKF81297-LY3154207 (orange), and D1R-SKF81297-LY3154207-UNC10062-BMS-A1 (magenta), each with the G_s_ protein removed. Notably, ICL3 exhibited significantly increased flexibility in the absence of PAMs. This dynamic behavior of ICL3 is visualized using the Cα atom of G246 (shown as spheres), which adopts a broader spatial distribution and frequently occupies the region normally filled by the bound G protein (grey surface). **d-f,** The conformational stabilization of Y331^7.53^ near the BMS-A1 binding site, the Cα atom positions of Y331^7.53^ are notably more confined in the triple-PAM-bound system, indicating a localized allosteric effect near the lower pocket.

The allosteric effects of these small-molecule PAMs may be obscured by the dominant influence of G protein interaction on receptor conformation. To isolate and evaluate the specific effects of BMS-A1 and UNC compounds from that of LY3154207 on D1R conformational dynamics in the absence of G protein, we performed MD simulations on three systems: D1R-SKF81297-only, D1R-SKF81297/LY3154207, and D1R- SKF81297/LY3154207/BMS-A1/UNC10062, after removing the G protein from the complex structure (Extended Data Table 9). In all simulations, the ligands remained stably bound to their respective sites and preserved their cryo-EM poses (Figs. 5c-f), consistent with their behavior in G protein-coupled simulations.

In the absence of G protein, we observed notable differences in local flexibility between PAM-bound and unbound systems. The intracellular loop 3 (ICL3) was significantly more dynamic without PAMs (Extended Data Fig. 17). By tracking the Cα atom of G246^ICL^^3^, a representative ICL3 residue, we found that in the SKF81297-only condition, G246^ICL^^3^ explored a broader space, frequently sampling conformations extending into the region that would be occupied by a bound G protein (Fig. 5c and Extended Data Fig. 17a). In contrast, the triple PAM-bound system showed a more compact distribution, suggesting that PAM binding stabilized ICL3 in a more open orientation even without the G protein. Moreover, our simulations indicated that the binding of BMS-A1 and UNC10062 stabilized the NPxxY motif in its active conformation (Figs. 5d-f, Extended Data Fig. 17b). Specifically, Y331^7.53^, a key residue within this conserved motif, remained highly flexible in the D1R-SKF81297-only and D1R- SKF81297/LY3154207 simulations but adopted a more stable conformation in the D1R-SKF81297/LY3154207/BMS-A1/UNC10062 system, resembling that observed in the G protein-bound state (Figs. 5d-f, Extended Data Fig. 17b). These findings support that BMS-A1 and UNC10062 promote a more ordered, potentially super-active receptor state by stabilizing TM7i and its associated microswitch motifs, and more broadly, by conferring structural rigidity to the intracellular regions of D1R in the absence of the G protein.

## Discussion

The allosteric modulation of GPCRs offers a promising avenue for developing safer and more effective treatments due to superior pharmacological profiles, such as enhanced receptor selectivity and finely-tuned pharmacological effects, compared to orthosteric ligands^6^. While single-site allosteric mechanisms have been extensively studied^6,34,39,43,50^, the mechanism of cooperative regulation of GPCR function by multiple modulators remains unexplored. The D1R has been proposed to harbor three spatially distinct PAM binding sites that function cooperatively^33^, providing an ideal system to investigate this multi-site allosteric modulation. Here, we provide the first comprehensive molecular elucidation of cooperative allosteric modulation in GPCRs. Combining medicinal chemistry, functional assays, cryo-EM analysis, and MD simulations, we show that allosteric modulators acting at three spatially distinct sites can synergize to enhance receptor activation: the ICL2 site occupied by LY3154207, the lower TM1-TM7 pocket occupied by BMS-A1, and the upper TM1-TM7 pocket occupied by the UNC compounds.

The vertically stacked arrangement of BMS-A1 and UNC compounds between TM1 and TM7, to our knowledge, represents a previously unreported configuration in GPCRs, highlighting how multiple modulators can coordinate to amplify signaling efficacy. Notably, the observation that cholesterol occupies a site similar to the UNC compounds suggests the possibility that these synthetic modulators may mimic lipid-based regulatory mechanisms. This observation aligns with broader evidence that cholesterol allosterically modulates various GPCRs, including the adenosine A2A receptor, oxytocin receptor, CC motif chemokine receptor 3, and D4R, indicating a potentially universal role for cholesterol in GPCR regulation^45,51–54^.

Mechanistically, our structural analyses reveal that these allosteric modulators enhance D1R activation through distinct yet complementary mechanisms. BMS-A1 promotes a more pronounced inward movement of TM7i, particularly the conserved NPxxY motif, while disrupting the T^1.46^-S^7.47^ interaction that maintains the inactive state. This facilitates the outward kink of TM6i necessary for G protein coupling^55–57^. Complementarily, UNC compounds stabilize the TM7 extracellular region in its active conformation by binding to the upper allosteric site. LY3154207 contributes by stabilizing the helical conformation of ICL2^39^. Together, these modulators establish a synergistic network that enhances receptor responsiveness to orthosteric agonists. It has also been suggested that LY3154207 disrupts the Na^+^-binding at D1R by compacting the Na^+^-binding pocket^39^, similar to the allosteric modulation of BMS986122 at a comparably located allosteric site in μOR^58,59^. Since BMS-A1 stabilizes the active TM7 conformation near the Na^+^ pocket, both BMS-A1 and LY3154207 may also facilitate D1R activation through disrupted Na^+^ binding.

Our analysis also elucidates the molecular basis of subtype selectivity of PAMs among DRs, a key consideration in drug development. S36^1.45^ facilitates BMS-A1 binding, whereas substitution by W53^1.45^ in D5R introduces steric hinderance and abolishes this interaction. By contrast, conservation of the upper pocket explains why the UNC scaffold exhibits activity at D5R. In D2-like receptors, the more significant divergence in both pockets underlies the selectivity of these compounds^24,25^. These insights establish a foundation for rational design of subtype-selective modulators with enhanced specificity and reduced off-target effects.

Currently, only a limited number of GPCRs, including β_2_-adrenergic receptor^60–62^, M_5_ muscarinic acetylcholine receptor^63,64^, FFAR2^43^, and metabotropic glutamate (mGlu) receptors^65–71^, have been reported to possess multiple allosteric sites. However, whether and how these spatially distinct sites cooperatively regulate a single GPCR remains poorly understood. The concept of simultaneously targeting multiple allosteric pockets within a single receptor as demonstrated in this study represents a paradigm shift applicable across the GPCR superfamily, offering remarkable opportunities for fine-tuning receptor function and developing combination therapies that exploit synergistic interactions. This approach may enable more precise control over receptor signaling and offer advantages over single-site modulators. In conclusion, our study reveals an unprecedented mechanism of cooperative allosteric modulation in D1R, where multiple spatially distinct sites act in concert to enhance receptor function. This discovery not only advances our understanding of GPCR signaling but also opens new avenues for the development of more effective therapies for neuropsychiatric disorders.

## Acknowledgments

The cryo-EM data of this study were collected at the Advanced Center for Electron Microscopy, Shanghai Institute of Materia Medica (SIMM). We thank all the staff at the center for their assistance in cryo-EM data collection. We also thank Shanghai Frontiers Science Center of Cellular Homeostasis and Human Diseases of Shanghai Jiao Tong University, School of Medicine for the technical support. This work was supported by the National Key R&D Program of China (2024YFA1307504 to Y.Z., 2022YFC2703105 to H.E.X.); the National Natural Science Foundation of China (32401002 to Y.Z., 32130022, 82121005, 82495184 to H.E.X.); the Young Elite Scientists Sponsorship Program by CAST (2023QNRC001 to Y.Z.); the CAS Strategic Priority Research Program (XDB37030103 to H.E.X.); Shanghai Municipal Science and Technology Major Project (2019SHZDZX02 to H.E.X.); Shanghai Municipal Science and Technology Major Project (H.E.X.); the Lingang Laboratory, Grant No. LG-GG-202204-01 (H.E.X.); the Natural Science Foundation of Shanghai, China (23ZR1475300 to Y.Z.); the Sailing Program of Shanghai Venus Project (23YF1456700 to Y.Z.); the “Shanghai Jiao Tong University 2030” Project-Category C (WH510363003/017 to Y.Z.); the Innovative Research Team of High-Level Local Universities in Shanghai (Y.Z.).

This research was supported in part by the Intramural Research Program of the National Institutes of Health (NIH). The findings and conclusions presented in this paper are those of the author(s) and do not necessarily reflect the views of the NIH or the U.S. Department of Health and Human Services. The contributions of the NIH author(s) are considered Works of the United States Government. Specific NIH funding was from the Intramural Research Programs of the National Institute of Neurological Disorders and Stroke (project ZIA-NS002263 to D.R.S.), the National Institute on Drug Abuse (project Z1A-DA000606 to L.S.), investigator initiated awards from the National Institute on Aging (R21 AG068833-01A1 to K.J.F.) and generous support provided by the Eshelman Innovation at the UNC Eshelman School of Pharmacy (project RX03812115 to K.J.F.). We acknowledge the University of North Carolina Department of Chemistry Mass Spectrometry Core Laboratory for their help with high-resolution mass spectrometry experiments to confirm compound identity. Research reported in this publication was supported by the Office of the Director, NIH under award number S10OD032476. The authors would like to thank Hannah Pearlstein, Natalie Asmus, and Eve Cohen for excellent technical assistance. Claude Sonnet 4 was used to refine the academic language and accuracy of the text.

## Author contributions

K.W. designed the constructs used for the purification of D1R-G_s_ complexes, purified the complexes, and prepared protein samples for the D1R-G_s_ complexes for cryo-EM data collection, determined the structure of D1R-G_s_ in complex with DA/LY3154207/BMS-A1/UNC9815, deposited all the structures, and prepared the draft of the manuscript and the figures; Y.W.X. performed cryo-EM grid preparation, data acquisition, structure determination of the D1R-G_s_ in complex with SKF81297/LY3154207/BMS-A1, and SKF81297/LY3154207/BMS-A1/UNC10062, built the models and refined the structures, and participated in manuscript preparation; Y.N.L. participated in cAMP assay and analysis, and figure preparation; W.H. participated in cryo-EM data collection; J.Z. supervised Y.N.L. in the cAMP assay; S.T.D., J.N.H. and K.D.L. performed the pharmacological assays and mutant receptor characterization; A.E.M., A.N.N. and R.B.F. supervised S.T.D. and J.N.H in the pharmacological assays, data analysis and figure preparation; F.W. and J.R. synthesized UNC9815 and UNC10062. K.J.F. designed UNC analogs and supervised F.W. and J.R. in synthetic chemistry; H.E.X. supervised K.W., Y.W.X., and W.H. in the cryo-EM studies; A.T., A.G., and B.X. carried out molecular modeling and simulation work and structural analysis; Y.Z., D.R.S, L.S., and Y.W. conceived of and supervised the project.

Y.Z. wrote the manuscript with inputs from all authors.

## Data availability

The cryo-EM density maps D1R-G_s_-Nb35 complexes bound to SKF81297/LY3154207/BMS-A1, SKF81297/LY3154207/BMS-A1/UNC10062, and Dopamine/LY3154207/BMS-A1/UNC9815 have been deposited in the Electron Microscopy Data Bank under accession numbers EMD-65916, EMD-65942 and EMD- 65949, respectively. The corresponding atomic coordinates have been deposited in the Worldwide Protein Data Bank (wwPDB) under accession numbers 9WEU, 9WG0 and 9WG6, respectively. All other data supporting the findings of this study are available in the main text and Extended Data Figures and Tables.

## Materials and Methods

### Compounds

Compounds UNC9815 and UNC10062 were synthesized in-house (see Extended Data Methods) at the UNC Eshelman School of Pharmacy (Chapel Hill, NC). BMS-A1 was a kind gift from Eli Lilly (Indianapolis, IN). LY3154207 was purchased from MedChemExpress (Monmouth Junction, NJ). Dopamine, SKF81297, and other fine chemicals were purchased from MilliporeSigma (St. Louis, MO) unless otherwise stated. DNA constructs for the BRET assays were kind gifts from Dr. Jonathan A. Javitch and Michel Bouvier. Receptor mutants were prepared at Bioinnovatise (Rockville, MD, USA), and all mutations were verified by DNA sequencing. Cell culture flasks, materials, and all assay plates were purchased from Thermo Fisher

Scientific (Waltham, MA) and Greiner Bio-One (Monroe, NC). Tissue culture media and related supplies were obtained from Invitrogen (Carlsbad, CA, USA) and Corning (Glendale, AZ, USA) unless stated otherwise.

### Construct design

The full-length gene coding sequence of human D1R was synthesized with no additional mutations or loop deletions added into the gene sequence. The N-terminal of D1R sequence was induced with FLAG tag followed by a fragment of β_2_AR N-terminal tail region (BN, hereafter) as fusion protein while the C-terminal was attached with an 8×His tag to facilitate the protein expression and purification^72^. The whole tag-attached D1R was then cloned into the standard pFastBac (Thermo Fisher) vector. The prolactin precursor sequence was inserted into the N-terminus before the FLAG tag as signaling peptide to assist cell membrane localization and increase expression of D1R. The NanoBiT tethering method was used to facilitate assembly of the D1R complexes, in which the C terminus of D1R was fused to the large part of NanoBiT (LgBiT), and the C terminus of G_β_ was fused to the small part of NanoBiT (SmBiT)^73^. A miniGαs format of G_αs_ with dominant negative mutations, G226A and A366S, was engineered (miniG_αs__DN) according to the previous study^36^. All the three G_s_ protein complex components, miniG_αs__DN, rat G_β1_ and bovine G_γ2_, were constructed into pFastbac vector separately.

### Expression, assemble and purification of D1R-Gs complexes

Before expression, the recombinant baculoviruses containing D1R, miniG_αs__DN, G_β1_ and G_γ2_ respectively were prepared using the Bac-to-Bac baculovirus expression system (Thermo Fisher). Cell cultures were grown to a density of 4 × 10^6^ cells/mL in ESF 921 serum-free medium (Expression Systems). For the expression of the D1R-G_s_ complexes, Sf9 cells were co-infected with the four types of baculoviruses prepared at 1:1:1:1 ratio. 48 hours after infection, the cultures were harvested by centrifugation at 1300 × g (Thermo Fisher, H12000) for 20 min and frozen at -80 °C for further usage.

For the purification of D1R-G_s_ in complex with SKF81297/LY3154207/BMS-A1, cell pellets from 1 L culture were thawed at room temperature and resuspended in buffer containing 20 mM HEPES, pH 7.2, 75 mM NaCl, 5 mM CaCl_2_, 5 mM MgCl_2_, 10% Glycerol, 0.3 mM TECP, protease inhibitor cocktail (Bimake, 1 mL/100 mL suspension). The cell pellets were dounced to homogeneity and subsequently added with 10 μM SKF81297 (Tocris) to induce complexes formation on cell membrane. Half an hour later, the cell suspension was further supplied with 20 μM LY3154207, treated with apyrase (25 mU/mL, NEB) and incubated at room temperature for another one hour. Finally, the cell membrane in suspension was solubilized directly by 0.5% (w/v) Lauryl Maltose Neopentyl Glycol (LMNG, Anatrace), 0.1% (w/v) cholesteryl hemisuccinate TRIS salt (CHS, Anatrace) and supplemented with 50 μM BMS-A1 and 10 μg/mL Nb35. The membrane was solubilized for 3 h at 4 °C before separation by ultracentrifugation. The sample was centrifuged at 100,000 × g (Ti45, Beckman) for 45 min. The isolated supernatant was then incubated for 2 h at 4 °C directly with FLAG resin (Smart-Lifesciences) pre-equilibrated with buffer containing 20 mM HEPES, pH 7.2, 100 mM NaCl. After binding, FLAG resin was washed with 20 column volumes of 20 mM HEPES, pH 7.2, 100 mM NaCl, 0.3 mM TCEP, 10 μM SKF81297, 10 μM LY3154207, 25 μM BMS-A1 0.01% (w/v) LMNG, 0.002% (w/v) CHS. The complex was eluted with 10 column volumes of 20 mM HEPES, pH 7.2, 100 mM NaCl, 0.3 mM TCEP, 10 μM SKF81297, 10 μM LY3154207, 25 μM BMS-A1, 0.01% (w/v) LMNG, 0.002% (w/v) CHS, 200 μg/μL FLAG peptide. The protein was then concentrated and loaded onto a Superose 6 Increase 10/300 column (GE Healthcare) pre-equilibrated with buffer containing 20 mM HEPES, pH 7.2, 100 mM NaCl, 0.3 mM TCEP, 0.00075% (w/v) LMNG, 0.00025% (w/v) glyco-diosgenin (GDN, Anatrace), 0.0002% (w/v) CHS and 10 μM SKF81297, 10 μM LY3154207, 25 μM BMS-A1. The fractions for the monomeric complex were collected and concentrated for electron microscopy experiments.

For the purification of D1R-G_s_ in complex with SKF81297/LY3154207/BMS- A1/UNC10062, all the purification processes are the same as the above with the following exceptions. The UNC10062 compound was added at the concentration of 50 μM with 10 μM SKF81297 (Tocris). In the later purification procedures, the buffer for washing and elution were all supplemented with 10 μM SKF81297, 10 μM LY3154207, 25 μM BMS-A1 and 25 μM UNC10062. The protein was loaded onto a Superdex 200 Increase 10/300 column (GE Healthcare) after concentration.

For the purification of D1R-G_s_ in complex with DA/LY3154207/BMS-A1/UNC9815, all the purification processes are the same as the above with the following exceptions. The UNC9815 compound was added at the concentration of 50 μM with 100 μM DA (MedChemExpress). In the later purification procedures, the buffer for washing and elution were all supplemented with 50 μM DA, 10 μM LY3154207, 25 μM BMS-A1 and 25 μM UNC10062. The protein was loaded onto a Superdex 200 Increase 10/300 column (GE Healthcare) after concentration.

### Cryo-EM grid preparation and data collection

For the preparation of cryo-EM grids, 3 μL of the purified complexes at 10.9 mg/mL for the D1R-G_s_ complex with SKF81297/LY3154207/BMS-A1, 16.0 mg/mL for the D1R-G_s_ complex with SKF81297/LY3154207/BMS-A1/UNC10062, 11.0 mg/mL for the D1R-G_s_ complex with DA/LY3154207/BMS-A1/UNC9815 were applied onto a glow-discharged holey carbon grid (Quantifoil R1.2/1.3). Grids were plunge-frozen in liquid ethane using Vitrobot Mark IV (Thermo Fisher Scientific). Frozen grids were transferred to liquid nitrogen and stored for data acquisition. Cryo-EM imaging was collected on a Titan Krios at 300 kV using Gatan K3 Summit detector and Falcon 4 direct electron detection device in the Cryo-Electron Microscopy Research Center, Shanghai Institute of Materia Medica, Chinese Academy of Sciences (Shanghai, China). A total of 6,244 movies for the D1R-G_s_ complex with SKF81297/LY3154207/BMS- A1 were collected on a Titan Krios equipped with a Gatan K3 direct electron detector with a pixel size of 0.824 Å. Images were taken at a dose rate of about 8.0 e/Å2/s with a defocus ranging from -1.0 to -2.0 μm using the EPU software (FEI Eindhoven, Netherlands). The total exposure time was 8 s, and 36 frames were recorded per micrograph. A total of 7,198 movies for the D1R-G_s_ complex with SKF81297/LY3154207/BMS-A1/UNC10062 and 8,517 movies for the D1R-G_s_ complex with DA/LY3154207/BMS-A1/UNC9815 were collected on a Titan Krios equipped with a Falcon 4 direct electron detection device at 300 kV. Images were taken with a pixel size of 0.73 Å, a defocus ranging from -1.0 to -2.0 μm using the EPU software (FEI Eindhoven, Netherlands) with total dose of 50 e/Å2/s over 2.5 s exposure on each EER format movie. Each movie was divided into 936 frames during motion correction.

### Image processing and map construction

For the D1R-G_s_ complex with DA/LY3154207/BMS-A1/UNC9815, all collected movie stacks were motion-corrected and dose-weighted in Relion4.0^74^. The contrast transfer function (CTF) parameters were estimated using the CTFFIND4 program within CryoSPARC v4^75^. Initial particle picking was performed with the Blob Picker on micrographs that had CTF parameters determined. The extracted particles underwent multiple rounds of 2D classification, yielding a subset of 390,276 particles. These were further refined through several iterations of ab initio reconstruction and heterogeneous refinement, ultimately converging on a well-defined particle subset. Subsequent heterogeneous refinement, non-uniform refinement, and local refinement were conducted in CryoSPARC v4^76^, resulting in a final reconstruction at a global resolution of 2.44 Å, derived from 209,112 particles.

### Model building and refinement

The structure of D1R -G_s_-SKF81297-LY3154207 (PDB:7LJC) was used as initial model for model rebuilding and refinement against the electron microscopy maps of D1R-G_s_ complexes. The initial models were docked into the electron microscopy density maps using ChimeraX followed by iterative manual adjustment and rebuilding in COOT^77^. Real space refinement and reciprocal space refinement were performed using Phenix programs^78^. Structure figures were prepared in ChimeraX-1.8.

### β-Arrestin Recruitment DiscoverX PathHunter Assay

Agonist-mediated D1R β-arrestin recruitment was determined using the DiscoveRx PathHunter complementation assay, as previously described by our laboratory^24,79^. Briefly, CHO-K1 cells stably expressing the D1R were maintained in Ham’s F12 supplemented with 10% fetal bovine serum, 100 U/mL penicillin, 100 μg/mL streptomycin, 800 μg/mL G418, and 300 μg/mL hygromycin, at 37 °C, 5% CO_2_, and 90% humidity. Cells were seeded in Cell Plating Media 2 (Eurofins DiscoverX, Fremont, CA) at a density of 2625 cells/well in 384-well black, clear-bottom plates. Following an 18- to 24-hour incubation, cells were treated with the indicated concentrations of compounds in phosphate-buffered saline (PBS) containing 0.2 mM sodium metabisulfite and then incubated at 37 °C for 90 minutes. The Tropix Gal-Screen buffer and substrate (Applied Biosystems, Bedford, MA) were added to cells according to the manufacturer’s recommendations and incubated for 30 min at room temperature. Luminescence signal was measured using a PHERAstar plate reader (BMG LABTECH, Cary, NC). Data were collected as RLUs and subsequently normalized to a percentage of the control luminescence seen with a maximum concentration of dopamine.

### Transient Transfections

HEK293 cells were grown in Dulbecco’s modified Eagle’s medium (DMEM) supplemented with 10% fetal bovine serum and maintained at 37 °C in a humidified incubator containing 5% CO_2_. The day before transfection, cells were plated in either 100 mm tissue culture dishes at 4 × 10^6^ cells/dish or 6-well plates at 1 × 10^6^ cells/well in serum-free DMEM. The cells were transfected with the indicated DNA constructs using polyethylenimine (PEI) as the transfection reagent at a ratio of 3:1 (μL PEI: μg DNA). 24 h after transfection, the media was changed to DMEM supplemented with 10% fetal bovine serum, and cells were used for experiments the next day.

### CAAX β-Arrestin Recruitment BRET Assay

HEK293 cells were transiently transfected with the indicated D1R constructs, GFP-CAAX (which is anchored in the plasma membrane^80^), and β-arrestin2-Rluc2. Cells were harvested with calcium-free Earle’s balanced salt solution (EBSS-), plated in 96- well white plates at 20,000 cells/well in Dulbecco’s phosphate-buffered saline (DPBS) and incubated at room temperature for 45 min. Cells were then incubated with 2 µM Prolume Purple (Nanolight Technology, Pinetop, AZ, USA) for 5 min, then stimulated with appropriate concentrations of test compounds for 5 minutes in < 3% DMSO. The BRET signal was determined by quantifying and calculating the ratio of the light emitted by GFP (515 nm) over Rluc2 (410 nm) using a PHERAstar FSX Microplate Reader (BMG Labtech, Cary, NC, USA).

### Direct β-Arrestin Recruitment BRET Assay

HEK293 cells were transiently transfected with the indicated D1R-Rluc8 constructs and β-arrestin2-mVenus as previously described^81^. Cells were harvested with EBSS-, plated in 96-well white plates at 20,000 cells/well in DPBS and incubated at room temperature for 45 min. Cells were incubated with 5 μM coelenterazine h (Nanolight Technology, Pinetop, AZ, USA) for 5 min and then stimulated with appropriate concentrations of test compounds for 5 minutes in < 3% DMSO. The BRET signal was determined by quantifying and calculating the ratio of the light emitted by mVenus (525 nm) over that emitted by Rluc8 (485 nm) using a PHERAstar FSX Microplate Reader (BMG Labtech, Cary, NC, USA).

### cAMP CAMYEL BRET Assay

HEK293 cells transiently expressing the indicated D1R constructs and the CAMYEL cAMP biosensor (yellow fluorescence protein-Epac-Rluc)^82^ were harvested with EBSS-, plated in 96-well white plates at 20,000 cells/well in DPBS and incubated at room temperature for 45 min. Cells were incubated with 5 μM coelenterazine h (Nanolight Technology, Pinetop, AZ, USA) for 5 min, then stimulated with appropriate concentrations of test compounds for 5 minutes in < 3% DMSO. The BRET signal was determined by quantifying and calculating the ratio of the light emitted by mVenus (525 nm) over that emitted by Rluc8 (485 nm) using a PHERAstar FSX Microplate Reader (BMG Labtech, Cary, NC, USA).

### Molecular modeling

Based on the cryo-EM structures reported in this study, the D1R-G_s_ complex models for the molecular dynamics (MD) simulations were constructed using MODELLER (version 10.6)^83^ to add key residues missing in the cryo-EM structures. Specifically, we added residue 20 of the N terminus, residues 169-184 of ECL2, residues 299-306 of ECL3, residues 241-263 of ICL3, and residues 346-348 of the Helix 8 (H8), using corresponding coordinates from a well equilibrated D1R-G_s_ model derived from our previous MD simulation study^39^. In addition, we ensured the formation of two disulfide bonds: C96^3.25^-C186^ECL2.50^ and C298^6.61^-C307^ECL^^3^ with constraints.

A total of 50 initial models were generated and assessed using the Discrete Optimized Protein Energy (DOPE) score^84^. The model with the lowest DOPE score, which also displayed loop orientations compatible with the membrane environment (i.e., not clashing with lipid molecules), was selected for the subsequent step.

### Molecular dynamics simulations

To build the MD simulation systems, the resulting D1R models generated with Modeller were processed using the Protein Preparation Wizard in Maestro (Schrödinger, version 2024.2). During this process, the hydrogen atoms were added, the protonation states of all titratable residues were assigned based on the pKa values predicted by PROPKA^85^, and Epik^86^ was used to determine the protonation and tautomeric states of the bound ligands in their bound conformations. We modeled residues D70^2.50^ and D120^3.49^ in their protonated (neutral) states, consistent with the current understanding of aminergic GPCRs in the active conformation. The nitrogen atoms of the orthosteric ligands (SKF and dopamine) were predicted to be protonated, carrying a positive charge. All histidine residues were kept neutral and C347 in H8 was modeled as palmitoylated.

The simulation systems were then constructed using the Desmond System Builder (Schrödinger, version 2024.2). Each D1R complex was embedded in a 1-palmitoyl-2- oleoyl-sn-glycero-3-phosphocholine (POPC) lipid bilayer and solvated with simple point charge (SPC) water model. The overall charge of each system was neutralized by adding appropriate counter ions, and 0.15 M NaCl was included to reproduce physiological ionic strength during MD simulations. The resulting system consists of approximately 165,000 atoms. The equilibrated system has approximate dimensions of ∼110 × 105 × 140 Å^3^.

MD simulations were performed with the Desmond MD engine (D. E. Shaw Research, New York, NY, USA) using OPLS4 force field. Ligand force field parameters for LY3154207 and UNC9815 were derived by quantum-mechanical optimization using the Schrödinger Force Field Builder (version 2024.2). Following our previously established GPCR simulation protocols^39^, each system was first minimized and equilibrated while applying harmonic positional restraints to ligand heavy atoms and protein backbone atoms (k = 1.0 kcal/mol/Å^2^). Simulations were carried out in the NPγT ensemble, with constant temperature (310 K) maintained using Langevin dynamics. Pressure was maintained at 1 atm using the hybrid Nose-Hoover Langevin piston method applied to an anisotropic flexible periodic cell.

During equilibration, a 1 fs integration time step was used and later increased to 2 fs for the production phase. Nonbonded interactions were calculated with a 12 Å cutoff. Harmonic restraints were gradually released during equilibration: first from the D1R loops, then from the transmembrane helices and bound ligands. Before production runs at 310 K, all restraints on the D1R were removed, while most α-carbon restraints on the G protein were retained, except for Gα residues 303-351 (α4-αG loop and α4 helix) and 383-394 (α5 helix). Long-timescale MD simulations were performed on V100x and A100 GPUs, with system coordinates saved every 600 ps. For each condition, at least three independent trajectories were generated with different random seeds. In total, 30 trajectories were obtained across all D1R systems (with and without PAMs), corresponding to an aggregated simulation time of ∼30 μs (Extended Data Table 9).

For the Gs protein-free D1R systems, we used the corresponding D1R-Gs complex models generated using Modeller, with the Gs protein removed, thereby ensuring the starting D1R conformation in the G-protein-free system matched that in the Gs-protein-bound system. This match allowed direct assessment of how the Gs-protein binding affects receptor dynamics. The resulting G protein-free systems were then processed, minimized, equilibrated, and subjected to the production runs using the protocols described above for the D1R-Gs complexes.

### Protein Interaction Analyzer (PIA)

PIA calculates pairwise distances between residues using either backbone Cα atoms or the centers of mass (COMs) of sidechain heavy atoms, and then determines distance differences between two conditions to assess conformational changes. The results are visualized as a heatmap, with the top-right and bottom-left halves representing Cα and sidechain-based distance differences, respectively. A key advantage of this method is that it does not rely on structural superposition, enabling robust analysis of rearrangement trends between two conditions^47^. In this study, pairwise distance differences of D1R binding-site residues were computed between inactive and active crystal structures, as well as between PAM-bound and unbound cryo-EM structures in the presence of orthosteric ligands (DA or SKF81297).

The binding-site analysis included following residues: 2.61, 2.64, 2.65, EL1.50, 3.28, 3.29, 3.32, 3.33, 3.36, 3.37, 3.40, 4.57, EL2.52, EL2.54, 5.38, 5.39, 5.42, 5.43, 5.46, 5.47, 5.50, 6.44, 6.48, 6.51, 6.52, 6.55, 6.56, 6.58, 6.59, 7.32, 7.35, 7.36, 7.39, 7.40, 7.42, and 7.43. Distance differences were plotted using a 1.0 Å cutoff. In the subsegment analysis, we defined the following subsegments in the transmembrane domain: TM1e (the extracellular section (e) of TM1, 1.30-1.36), TM1m (the middle section (m) of TM1, residues 1.37-1.45),TM1i (the intracellular section (i) of TM1, residues 1.46-1.59), TM2i (residues 2.38-2.51), TM2m (residues 2.52-2.60), TM2e (residues 2.61-2.66), TM3e (residues 3.22-3.35), TM3m (residues 3.36-3.40), TM3i (residues 3.41-3.55), TM4i (residues 4.39-4.49), TM4m (residues 4.50-4.55), TM4e (residues 4.56-4.62), TM5e (residues 5.36-5.45), TM5m (residues 5.46-5.50), TM5i (residues 5.51-5.63), TM6i (residues 6.30-6.43), TM6m (residues 6.44-6.48), TM6e (residues 6.49-6.60), TM7e (residues 7.32-7.43), TM7i (residues 7.44-7.54).

### Pairwise RMSD calculations

Pairwise root mean square deviation (RMSD) distributions were calculated for the PAMs (BMS-A1, UNC10062, and UNC9815) using VMD^87^. For each trajectory, the first 100 frames were discarded. For each pair of frames, the frames were superimposed on the Cα atoms of the PAM binding-site residues (1.38, 1.39, 1.41, 1.42, 1.45, 1.48, 1.49, 7.36, 7.37, 7.40, 7.41, 7.44, 7.48, 7.51, 7.55, and 8.54), after which the RMSD was calculated using ligand heavy atoms. RMSD values computed between all frame pairs yielded pairwise RMSD distributions that quantify the stability and conformational dynamics of PAMs within their binding sites.

### Statistical analysis

The data were analyzed using GraphPad Prism 10.2.3 (GraphPad Software, Inc., La Jolla, CA). All results are normalized to dopamine control. Potencies, fold-shifts, and maximum efficacies are displayed as mean ± SEM. Statistical significance was determined using a two-tailed *t*-test when two groups were compared and using a one-way analysis of variance (ANOVA) when multiple groups were compared, with p < 0.05 used as the cutoff for statistical significance (pEC_50_ values were used for the statistical analyses).

### Synthesis of UNC9815 and UNC10062

#### (1) General Information

All reagents were used as received from commercial sources (Sigma Aldrich, TCI America and ThermoFisher). 4-(*tert*-Butyl) cyclohexanone **S1** and methyl 2-(4- hydroxyphenyl) acetate **S6** were purchased from Ambeed Scientific. The ^1^H and ^13^C

NMR spectra were recorded on a 400 MHz Bruker Avance spectrometer equipped with a broadband observe probe and a 500 MHz Bruker AVIII spectrometer equipped with a dual cryoprobe, respectively. Chemical shifts are reported in parts per million and were referenced to residual proton solvent signals. Microwave syntheses were conducted in a Biotage Initiator constant temperature microwave synthesizer. Flash column chromatography separations were performed by the use of the Teledyne Isco CombiFlash Rf using RediSep Rf silica gel columns eluting with hexanes/ethyl acetate (0 to 100%). Thin-layer chromatography (TLC) was performed on Analtech UNIPLATE silica gel GHLF plates (gypsum inorganic hard layer with fluorescence). TLC plates were developed using iodine vapor, potassium permanganate stain or ceric ammonium molybdate stain, as required. Compound purity was measured on the basis of peak integration (area under the curve) from UV/Vis absorbance (at 214 nm) on a Waters Acquity ultra performance LC system with a photodiode array detector coupled to a Waters Acquity single quadrupole detector mass spectrometer with an ESI source. Compound identity was determined on the basis of HRMS analysis. HRMS samples were analyzed at the UNC Department of Chemistry Mass Spectrometry Core Laboratory using a Q Exactive HF-X mass spectrometer.

#### (2) Synthetic Procedures and Characterization of UNC9815

2-(4-(*tert*-Butyl) cyclohexylidene)-2-cyanoacetamide (**S3**). To a solution of 4-(*tert*- butyl) cyclohexanone **S1** (500 mg, 3.24 mmol) and 2-cyanoacetamide **S2** (273 mg, 3.24 mmol, 1.0 equiv) in toluene (50 mL) was added ammonium acetate (500 mg, 3.24 mmol, 1.0 equiv) and acetic acid (185 µL, 3.24 mmol, 1.0 equiv) at room temperature. The reaction was heated to reflux and water removed azeotropically using a Dean–Stark trap. After complete evolution of water, the reaction was cooled to room temperature. Water (50 mL) and ethyl acetate (25 mL) were added and the layers separated. The organic layer was dried over sodium sulphate and the solvent evaporated under vacuum. The residue was purified by flash chromatography (40% ethyl acetate in hexanes) to give **S3** (475 mg, 2.16 mmol, 67% yield) as a white solid. ^1^H-NMR (500 MHz, CDCl_3_): δ 0.87 (d, *J* = 1.3 Hz, 9H), 1.17 – 1.38 (m, 3H), 1.97 – 2.04 (m, 2H), 2.04 – 2.11 (m, 1H), 2.23 – 2.34 (m, 1H), 3.02 (ddd, *J* = 13.9, 3.7, 2.3 Hz, 1H), 3.94 – 4.03 (m, 1H), 6.00 (d, *J* = 128.1 Hz, 2H).

2-Amino-6-(*tert*-butyl)-4,5,6,7-tetrahydrobenzo[b]thiophene-3-carboxamide (**S4**). To a solution of **S3** (400 mg, 2.16 mmol) in ethanol (10 mL) was added sulphur (76 mg, 2.37 mmol, 1.1 equiv) and morpholine (186 µL, 2.16 mmol, 1.0 equiv) at room temperature. The reaction was heated to 50 °C and stirred for 5 h. After completion, the solvent was evaporated under vacuum. The residue was purified by flash chromatography (40% ethyl acetate in hexanes) to give **S4** (489 mg, 1.94 mmol, 90% yield) as a white solid. ^1^H-NMR (500 MHz, DMSO-*d*_6_): δ 0.91 (s, 9H), 1.13 – 1.27 (m, 1H), 1.36 – 1.45 (m, 1H), 1.93 (d, *J* = 9.4 Hz, 1H), 2.21 – 2.30 (m, 1H), 2.47 (dd, *J* = 15.7, 4.9 Hz, 1H), 2.60 (d, *J* = 13.2 Hz, 1H), 2.69 (dd, *J* = 15.7, 4.9 Hz, 1H), 6.49 (s, 2H), 6.92 (s, 2H). These data match those previously reported^24^.

2-Amino-6-(*tert*-butyl) benzo[b]thiophene-3-carboxamide (**S5**). To a solution of **S4** (489 mg, 1.94 mmol) in 1,4-dioxane (10 mL) was added chloranil (953 mg, 3.88 mmol, 2.0 equiv) and the reaction was heated to reflux for 17 hours while monitoring the progress of the reaction using UPLC-MS. The solvent was evaporated under vacuum and the residue partitioned between dichloromethane and water. The biphasic mixture was filtered through celite and the layers separated. The organic layer was dried over sodium sulfate and evaporated under vacuum. The residue was purified by flash chromatography (15% ethyl acetate in hexanes) to give **S5** (341 mg, 1.37 mmol, 71% yield) as a pale pink solid. ^1^H-NMR (400 MHz, CDCl_3_): δ 1.35 (s, 9H), 5.65 (s, 2H), 6.55 (s, 2H), 7.37 (dd, *J* = 8.5, 1.9 Hz, 1H), 7.51 (dd, *J* = 8.5, 0.6 Hz, 1H), 7.57 (dd, *J*= 1.9, 0.6 Hz, 1H); ^13^C-NMR (100 MHz, CDCl_3_): δ 31.6, 34.8, 101.5, 118.98, 119.02, 123.5, 129.9, 134.4, 145.6, 162.6, 168.6.

*N*-(6-(*tert*-Butyl)-3-carbamoylbenzo[b]thiophen-2-yl) isonicotinamide (**UNC9815**). To a solution of isonicotinic acid (12.0 mg, 0.10 mmol) and catalytic amount (10 µL) of DMF in anhydrous dichloromethane (2 mL) was added oxalyl chloride (13 µL, 0.15 mmol, 1.5 equiv) and the reaction stirred for 2 h at room temperature. The solvent was evaporated under vacuum, the residue was dissolved in anhydrous dichloromethane (2 mL), **S5** (25.0 mg, 0.10 mmol, 1.0 equiv) and pyridine (16 µL, 0.20 mmol, 2.0 equiv) were added, and the reaction was stirred for 2 h at room temperature. Progress of the reaction was monitored by TLC and, upon completion, the solvent was removed under vacuum. The residue was extracted with ethyl acetate (25 mL), filtered and again the solvent was removed under vacuum. Finally, the residue was purified by flash chromatography (5% methanol in dichloromethane) to afford *N*-(6-(*tert*-butyl)-3-carbamoylbenzo[b]thiophen-2-yl) isonicotinamide **UNC9815** (25.0 mg, 0.07 mmol, 69% yield) as an off-white solid. TLC (DCM:MeOH, 95:5, v/v): *Rf* = 0.5; ^1^H NMR (400 MHz, DMSO-*d*_6_) δ 1.37 (s, 9H), 7.55 (dd, *J* = 8.7, 1.9 Hz, 1H), 7.84 – 7.87 (m, 2H), 7.91 (s, 2H), 7.99 (d, *J* = 8.6 Hz, 1H), 8.01 (d, *J* = 1.8 Hz, 1H), 8.88 – 8.92 (m, 2H), 13.43 (s, 1H); ^13^C NMR (100 MHz, DMSO-*d*_6_) δ 31.3, 34.6, 111.2, 118.6, 120.9, 121.7, 123.4, 130.7, 133.9, 139.2, 146.1, 146.8, 151.0, 162.0, 167.6; UPLC purity = 99%; HRMS (m/z): [M+H]^+^ calcd for C_19_H_20_N_3_O_2_S, 354.1271; found, 354.1268.

#### (3) Synthetic Procedures and Characterization of UNC10062

Methyl 2-(4-(benzyloxy) phenyl) acetate (**S7**). According to the protocol of Qiang Liu and co-workers^88^. A solution of methyl 2-(4-hydroxyphenyl) acetate **S6** (1.53 g, 9.19 mmol), benzyl bromide (1.57 g, 9.19 mmol, 1.0 equiv) and potassium carbonate (3.81 g, 27.58 mmol, 3.0 equiv) in DMF (20 mL) was stirred at room temperature for 15 h. Water (50 mL) was added, and the reaction extracted with EtOAc (3 × 50 mL). The combined organic layers were washed with brine, dried with Na_2_SO_4_ and concentrated under reduced pressure. The mixture was further purified flash chromatography to give **S7** (2.01 g, 7.84 mmol, 85% yield). ^1^H-NMR (400 MHz, CDCl_3_): δ 3.57 (s, 2H), 3.69 (s, 3H), 5.05 (s, 2H), 6.91 – 6.96 (m, 2H), 7.18 – 7.23 (m, 2H), 7.29 – 7.46 (m, 5H).

Methyl 2-(4-(benzyloxy) phenyl)-2-methylpropanoate (**S8**). According to the protocol of Qiang Liu and co-workers^88^. A solution of **S7** (2.00 g, 7.80 mmol) in THF (20 mL) was slowly added to a slurry of sodium hydride (935.7 mg, 60 wt %, 23.39 mmol, 3.0 equiv) in THF (20 mL) at 0 °C, then stirred at room temperature for 1 h. The reaction was cooled to 0 °C, and iodomethane (3.32 g, 23.39 mmol, 3.0 equiv) was added. The mixture was stirred at room temperature for 15 h. Water (50 mL) was added, and the reaction extracted with EtOAc. The combined organic layers were dried with Na_2_SO_4_ and concentrated under reduced pressure. The mixture was further purified by flash chromatography to give **S8** (0.6489 g, 2.282 mmol, 29% yield). ^1^H-NMR (400 MHz, CDCl_3_): δ 1.55 (s, 6H), 3.63 (s, 3H), 5.03 (s, 2H), 6.87 – 6.96 (m, 2H), 7.20 – 7.27 (m, 2H), 7.28 – 7.43 (m, 5H).

Methyl 2-methyl-2-(4-oxocyclohexyl) propanoate (**S9**). According to the protocol of Qiang Liu and co-workers^88^. To a solution of methyl 2-(4-(benzyloxy) phenyl)-2- methylpropanoate **S8** (0.63 g, 2.22 mmol) in 1,2-dichloroethane (10 mL) in a Parr reactor 25 mL glass insert was added Pd/Al_2_O_3_ (0.14 g, 5 wt%). The 25 mL glass insert was placed in a 150 mL stainless steel Parr reactor under air. The Parr reactor was pressurized and depressurized with hydrogen gas three times before the indicated pressure (5 bar) was set. The reaction was stirred and heated at 60 °C for 24 h. Filtration of the catalyst through a pad of celite and concentration of the filtrate gave crude product which was further purified by flash chromatography to give **S9** (0.24 g, 1.19 mmol, 54% yield).^1^H-NMR (400 MHz, CDCl_3_) δ 1.17 (s, 6H), 1.49 (qd, *J* = 13.0, 5.1 Hz, 2H), 1.87 – 2.00 (m, 2H), 2.09 (ddd, *J* = 12.1, 9.2, 3.0 Hz, 1H), 2.29 – 2.47 (m, 4H), 3.69 (s, 3H). These data are in agreement with those previously reported^89^.

Methyl 2-(2-amino-3-carbamoyl-4,5,6,7-tetrahydrobenzo[*b*]thiophen-5-yl)-2- methylpropanoate (**S10**). A solution of **S9** (0.38 g, 1.90 mmol), 2-cyanoacetamide (239.3 mg, 2.85 mmol, 1.5 equiv), sulfur (60.82 mg, 1.90 mmol, 1.0 equiv), and morpholine (247.9 mg, 2.85 mmol, 1.5 equiv) in EtOH (3 mL) was heated under microwave irradiation at 120 °C for 30 min. The reaction was concentrated under reduced pressure and the residue purified by flash chromatography to give **S10** (0.3394 g, 1.145 mmol, 60% yield). ^1^H-NMR (400 MHz, CD_3_CN): δ 1.14 (s, 3H), 1.18 (s, 3H), 1.34 (qd, *J* = 12.5, 5.2 Hz, 1H), 1.86 (ddd, *J* = 12.9, 5.4, 1.9 Hz, 1H), 1.96 – 2.05 (m, 1H), 2.31 – 2.48 (m, 2H), 2.51 – 2.62 (m, 1H), 2.76 (dd, *J* = 15.7, 5.3 Hz, 1H), 3.63 (s, 3H), 5.61 (s, 2H), 6.38 (s, 2H); ^13^C NMR (126 MHz, CD_3_CN): δ 22.3, 22.5, 25.6, 26.6, 28.2, 43.3, 45.9, 52.2, 108.1, 130.5, 161.6, 16.9, 178.5; UPLC purity = 99%; HRMS (*m/z*) [M + H]^+^: calcd for C_14_H_21_N_2_O_3_S, 297.1267; found, 297.1268.

Methyl 2-(3-carbamoyl-2-(isonicotinamido)-4,5,6,7-tetrahydrobenzo[*b*]thiophen-6- yl)-2-methylpropanoate (**UNC10062**). To a solution of **S10** (0.54 g, 1.81 mmol) and triethylamine (549.5 mg, 5.43 mmol, 3.0 equiv) in DCM (10 mL) at 0 °C was slowly added isonicotinoyl chloride hydrochloride (526.8 mg, 2.72 mmol, 1.5 equiv). The reaction was stirred for 4 h, quenched by the addition of saturated aqueous NaHCO_3_ (5 mL), and extracted with DCM (3 × 20 mL). The combined layers were dried with Na_2_SO_4_, and purified by flash chromatography to give **UNC10062** (234.1 mg, 0.58 mmol, 32% yield). ^1^H-NMR (400 MHz, CDCl_3_): δ 1.23 (s, 3H), 1.24 (s, 3H), 1.49 (qd, *J* = 12.5, 5.2 Hz, 1H), 1.98 (dd, *J* = 12.9, 3.4 Hz, 1H), 2.07 – 2.17 (m, 1H), 2.50 – 2.63 (m, 1H), 2.69 (dd, *J* = 15.8, 4.8 Hz, 2H), 2.95 (dd, *J* = 14.6, 5.1 Hz, 1H), 3.71 (s, 3H), 5.94 (s, 2H), 7.82 – 7.88 (m, 2H), 8.83 (d, *J* = 5.8 Hz, 2H), 13.40 (s, 1H); ^13^C-NMR (101 MHz, CDCl_3_): δ 21.7, 22.5, 24.8, 25.8, 27.2, 41.9, 45.1, 51.9, 113.6, 121.1, 127.6, 128.5, 139.9, 146.7, 150.5, 161.4, 168.4, 177.7; UPLC purity = 97%; HRMS (*m/z*) [M + H]^+^: calcd for C_20_H_24_N_3_O_4_S, 402.1482; found, 402.1481.

